# Mucin granules are degraded in the autophagosome-lysosome pathway as a means of resolving airway mucous cell metaplasia

**DOI:** 10.1101/2020.06.02.130534

**Authors:** JM Sweeter, K Kudrna, K Hunt, P Thomes, BF Dickey, SL Brody, JD Dickinson

## Abstract

Exacerbations of muco-obstructive airway diseases such as COPD and asthma are associated with epithelial changes termed mucous cell metaplasia (MCM). The molecular pathways triggering MCM have been identified; however, the factors that regulate resolution are less well understood. We hypothesized that the autophagosome-lysosome pathway is required for resolution of MCM by degrading cytoplasmic mucins. We found increased intracellular levels of Muc5ac and Muc5b in autophagy-deficient mice. This difference was not due to defective mucin secretion. Instead, we found that Lamp1-labeled lysosomes surrounded mucin granules of mucous cells indicating that granules were being degraded. Using a model of resolution of mucous cell metaplasia in mice, we found increased lysosomal proteolytic activity that peaked in the days after inflammation. Autophagy-deficient mice had persistent accumulation of mucin granules that failed to decline due to reduced mucin degradation. We applied these findings *in vitro* to human airway epithelial cells (AECs). Activation of autophagy by mTOR inhibition led to degradation of mucin granules in AECs. Our findings indicate that during peak and resolution phases of MCM, mucin granules can be degraded by autophagy. The addition of mucin degradation to the existing paradigm of production and secretion may more fully explain how the secretory cells handle excess amounts of cytoplasmic mucin and offers a therapeutic target to speed resolution of MCM in airway disease exacerbations.

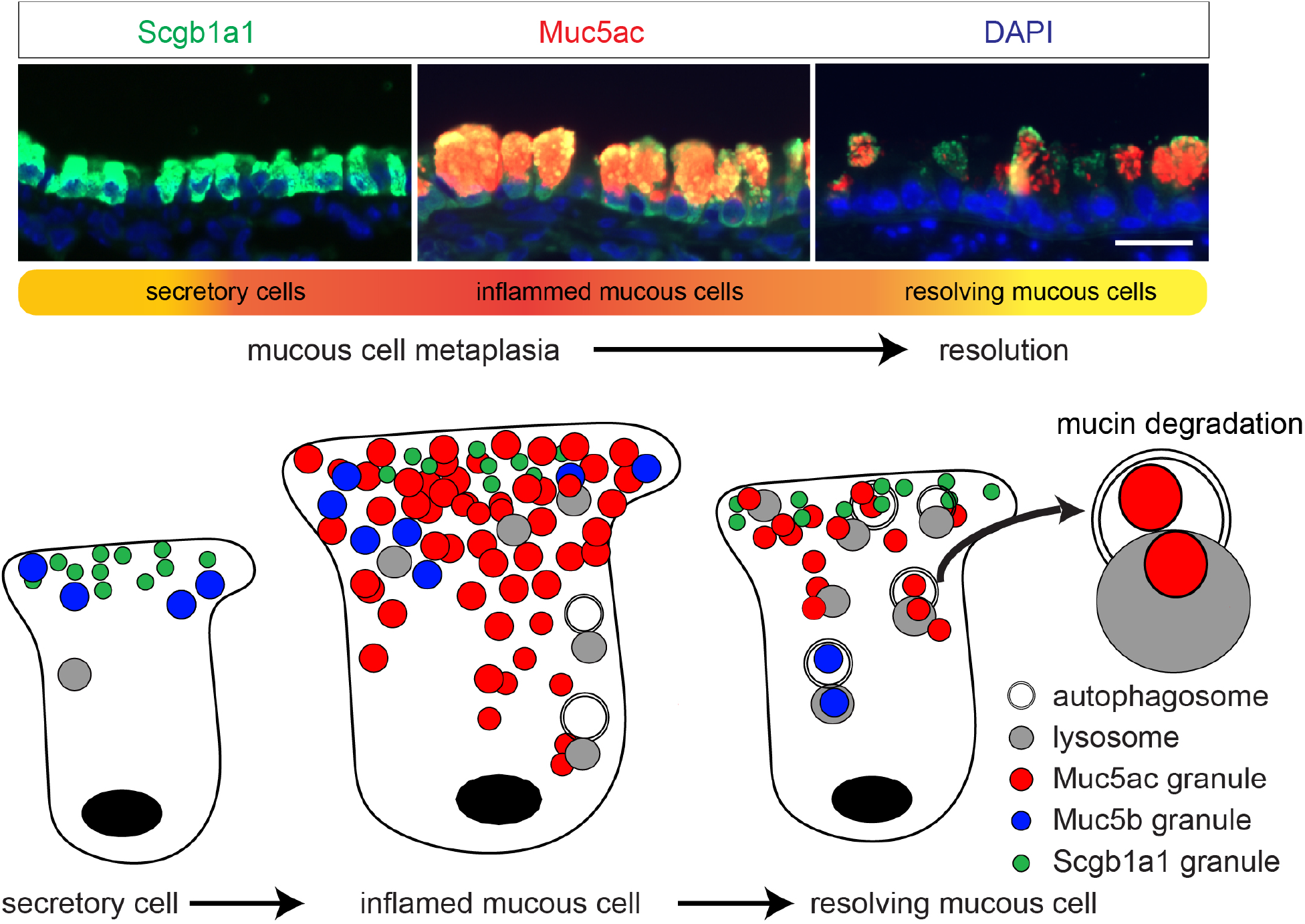

## Introduction

Mammalian cells utilize two primary degradation systems to recycle proteins: the proteasome and the autophagosome-lysosome pathways. While there is some overlap, these pathways are largely independently regulated and serve different functions(1). These protein degradation systems are necessary for the cell to balance metabolic demands by recycling proteins to amino acids for new synthesis. The autophagosome-lysosome pathway is highly conserved across species(2, 3). It can be utilized in bulk protein breakdown (macro-autophagy or referred to as autophagy) in response to nutrient demands for new amino acids. Cells can also utilize chaperone-mediated autophagy that utilizes a specific amino acid motif targeted by cytoplasmic chaperones that bypass the autophagosome and directly insert proteins in the lysosome for degradation. Finally, cells can utilize selective autophagy by targeting proteins, protein aggregates, and organelles including mitochondria(4, 5) and even cilia components(6) to the autophagosome for degradation in the lysosome. We(7, 8), and others(9–13), have observed that autophagy markers are increased in models of human airway disease, and in asthma and COPD airways. This led us to infer that autophagy played a key role in the response to airway inflammation in the epithelium.

Mucociliary clearance is a vital feature of innate immunity(14, 15). There are two primary secretory mucins in the murine and human airways, MUC5B and MUC5AC, that provide the biophysical properties of mucous gel layer(16, 17). MUC5B is the homeostatic mucin tonically produced and secreted throughout the airway to facilitate mucocilary clearance(18). In contrast, MUC5AC production and secretion is highly inducible in response to airway inflammation(19, 20). Mucins are large proteins, glycosylated in the Golgi and then packaged in granules for apical transport to the plasma membrane where they await secretion to the airway lumen. Mucin proteins can undergo baseline or stimulated secretion(21–26) and rely on distinct protein sorting receptors to prepare granules for release by either method(27).

In muco-obstructive airway diseases such as asthma, chronic obstructive pulmonary disease (COPD), and cystic fibrosis, excessive MUC5AC production leads to extensive changes of the airway epithelium, termed mucous cell metaplasia, marked by enlargement of mucous cells with increased intracellular staining of MUC5AC. Airway inflammation provides stimuli through the action of ATP for enhanced mucin secretion into the airway leading to airway obstruction and loss of lung function(19, 28). The factors that regulating the development of mucous cell metaplasia in muco-obstructive airway disease have been extensively characterized in mouse and human airway models. In the mouse, secretory cells marked by cytoplasmic Secretoglobin 1a1, Muc5b, along with proteins involved in the regulated exocytic machinery transition from a club cell appearance to goblet cell morphology with cytoplasm packed with Muc5ac granules(21). Depending on the model, 50-300-fold inductions of Muc5ac have been observed(8, 20, 29, 30). The sheer magnitude of Muc5ac production leads to ER stress responses(31) and activated unfold protein responses(32) along with epithelial ROS(8). A number of inflammatory and allergen factors have been identified that initiate mucous cell metaplasia of the airway epithelium. These include Type 2 inflammatory cytokines such as IL-13(33–35), growth and development factors such as NOTCH(36–38) and EGF(39), and respiratory viruses such as rhinovirus(40). The transcriptional activity of SPDEF(41, 42), plays a significant role downstream of many of the signals to induce mucous cell metaplasia. What has not been studied is how the airway secretory cell normalizes as airway inflammation abates. We considered that during resolution mucin granules may continue to be secreted, or alternatively, degraded.

Here, we propose that the autophagy-lysosome pathway is utilized to accelerate resolution by breaking down cytoplasmic mucins. In support of this we show that mice globally deficient in the autophagy regulatory protein, Atg16L1, have reduced proteolytic function. Additionally, they have increased accumulation of cytoplasmic mucin granules in models of type 2 airway inflammation particularly during resolution of mucous cell metaplasia. We also found lysosomal markers closely approximate and engulf mucin granules. Finally, we demonstrate that induction of autophagy through mTOR inhibition led to MUC5AC degradation in human AECs derived from non-diseased and COPD derived sources.

## Results

### Atg16L1^HM^ mice have increased airway Muc5ac and Muc5b levels during mucous cell metaplasia

We previously showed that in human AECs cultured with IL-13, there was increased cytoplasmic positive MUC5AC granules when autophagy protein, ATG5 was depleted by shRNA(7). In addition, we found that that LC3 positive autophagosome levels were increased as measured by LC3 II accumulation during flux assay in human AECs challenged with type 2 cytokines IL-13 and IL-4(7, 8). ATG5-12 and ATG16L1 form a complex that facilitates autophagosome maturation. We, therefore utilized mice globally deficient in Atg16L1 (Atg16L1^HM^) which have reduced degradation of autophagy target Lc3 II at baseline in the lungs (**Supplemental Figure 1 A,B**) consistent with previous findings in the mouse ilium(43). To explore the effect of deficient autophagy mediated degradation during states of significant mucin production and intracellular accumulation, we stimulated wildtype (WT) or Atg16L1^HM^ mice with 3 intra-nasal IL-33 challenges over 7 days. Atg16L1^HM^ mice had increased cytoplasmic Muc5ac and to a lesser extent Muc5b positive granules by immunostaining compared to WT mice (**Figure 1 A,-C**). A similar finding was seen by immunoblot from mouse lung homogenates with higher levels of Muc5ac in Atg16L1^HM^ mice (**Figure 1 D and E**). These findings indicate the mice globally deficient in protein degradation by autophagy, have increased intracellular accumulation of Muc5ac and trend toward higher Muc5b.

**Figure 1:**
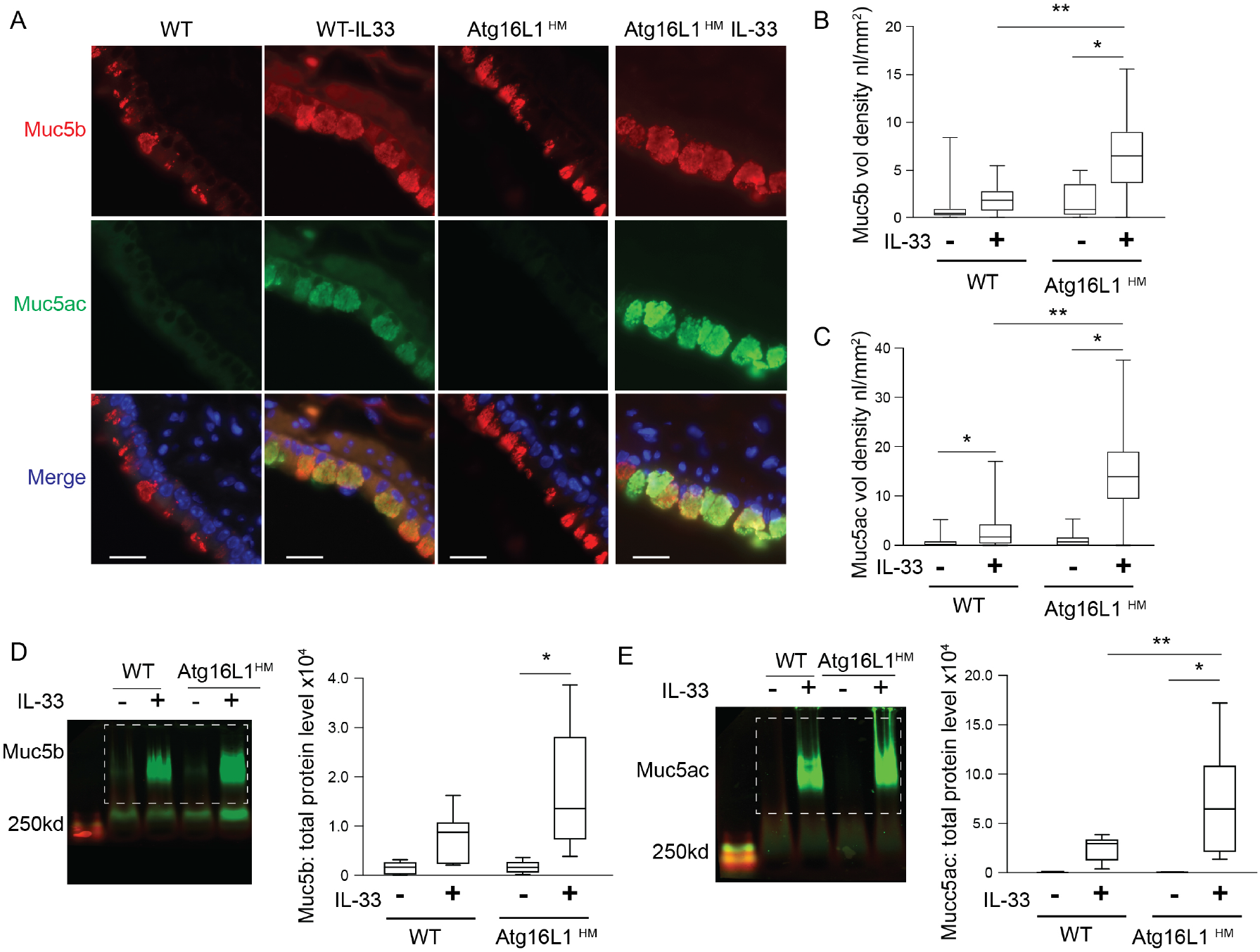
Atg16L1 deficient mice have increased cytoplasmic mucins with IL-33 challenge. (**A**) Representative images of immunostaining of mouse anti-Muc5b (red) and Muc5ac by lectin UEA-1 (green) are shown. Nuclei are countered stained with DAPI in blue. Scale bar equals 20 microns. Quantification of Muc5b (**B**) and Muc5ac (**C**) airways volume density (nl/mm2 (n=6 mice per group). (**D-G**) Representative immunoblots of lung homogenates of Muc5b and Muc5ac from Atg16L1^HM^ or WT mice with corresponding quantification normalized to total protein level (n=7 mice per group). Dashed box denotes area of mucin quantification. Graphs show box and whisker blots with min and max values on whiskers and quartile box with median value. Significant difference by ANOVA with Tukey's post-hoc comparisons are noted by * for IL-33 treatment difference or ** for mouse genotype differences.

### Atg16L1 deficient mice have a normal secretory response

To test if the accumulation of airway mucins in Atg16L1^HM^ mice was related to dysfunctional mucin secretion, we nebulized ATP immediately before euthanasia in OVA sensitized and challenged mice. As was the case with IL-33 challenge, inflammation by OVA led to the increased Muc5ac in the lungs of mice deficient of Atg16L1 and a trend toward increased Muc5b (**Figure 2A and B**). We examined by immunofluorescence the intracellular mucin granule content. ATP led to the expected decrease of intracellular Muc5b and Muc5ac granules in both the WT and Atg16L1^HM^ mice (**Figure 2 C-E**). Many of the individual secretory cells after exposure to ATP lost the feature of tightly organized circular mucin granules. Instead there was irregular mucin staining and reduced intensity suggesting that secretory cells had released many of the preformed granules in response to ATP. In the airways, we found examples of complete occlusion by mucous plugs primarily consisting of Muc5ac in both the WT and Atg16L1^HM^ mice (**Figure 2F**). These data indicate that the accumulation of airway mucins was not due to defects in ATP-mediated secretion. Secretory cells also rely on baseline mucin secretion with distinct exocytic machinery(22, 27, 44). As Muc5b is the primary mucin produced and secreted in the naïve mouse airway, we measured Muc5b levels in whole lung homogenates as a marker of baseline mucin secretion in WT and Atg16L1^HM^ mice. We found no significant difference in Muc5b levels in the lungs from autophagy deficient mice nor in Muc5b intracellular staining in mouse AEC from WT and Atg16L1^HM^ mice differentiated under air liquid interface (ALI) conditions for 14 days (**Figure 3 A,B**). There was also no significant difference in the levels of secretory cell marker, Scgb1a1, obtained during a time course of ALI differentiation indicating a normal differentiation program in Atg16L1 deficient AECs (**Figure 3C**). Therefore, we deduced that both baseline and ATP-stimulated secretion pathways remain intact and secretory cells differentiate normally in autophagy deficient mice. We therefore explored a degradation fate for cytoplasmic mucins in the airway epithelium.

**Figure 2:**
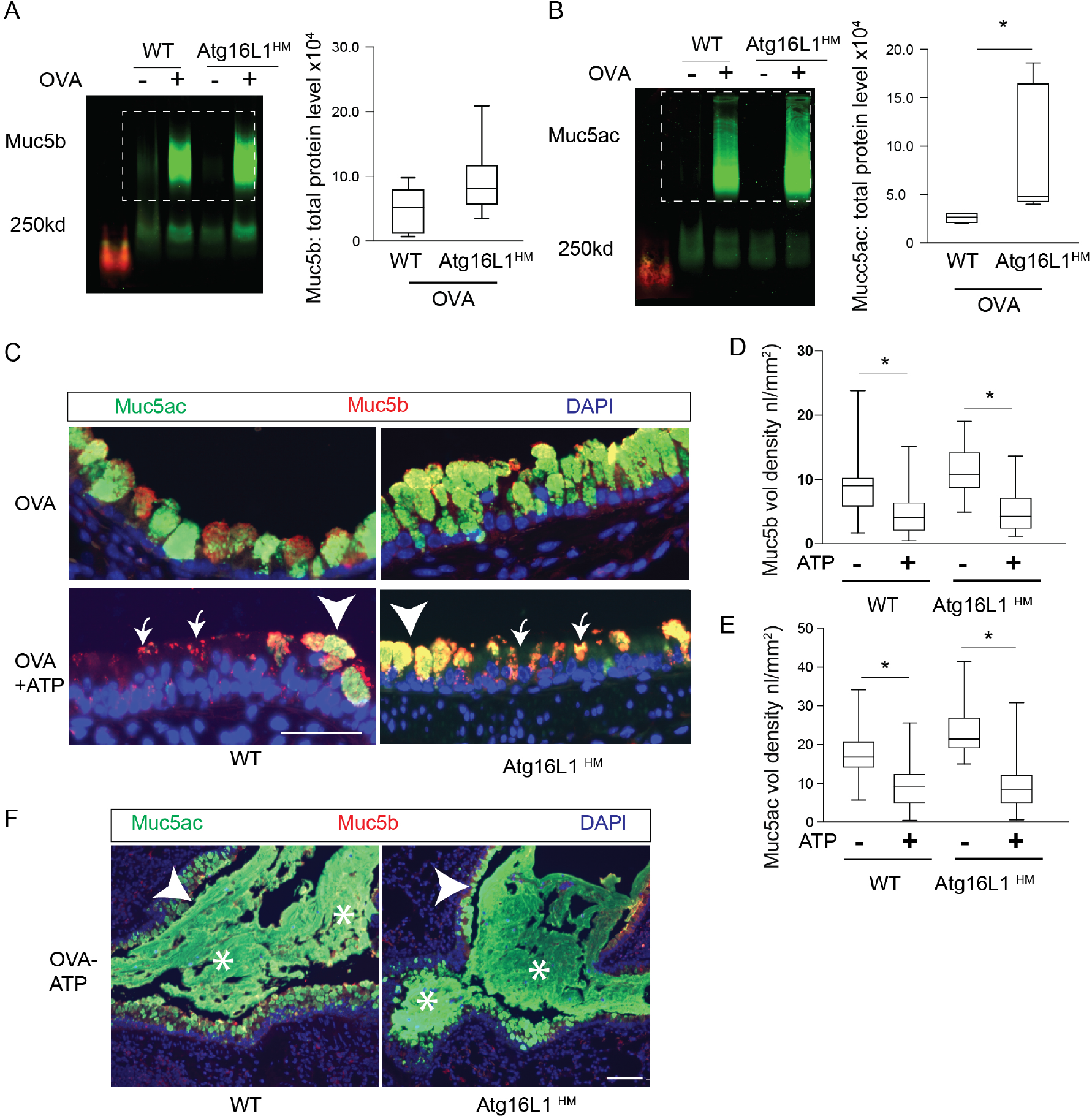
A,B) Atg16L1 deficient mice have normal mucin secretory response. Representative immunoblots of lung homogenates of Muc5b and Muc5ac from OVA challenged WT or Atg16L1^HM^ mice with corresponding quantification normalized to total protein level (n=6 mice per group) (**C**) Representative images of immunostaining of mouse anti-Muc5b and Muc5ac by lectin UEA1 in OVA challenged Atg16L1 ^HM^ and WT mice ± ATP nebulization with corresponding quantification of mucin levels (nl/mm^2^) for Muc5b (**D**) and Muc5ac (**E**) (n=6 mice per group). Arrows point to mucous cells that have recently emptied cytoplasmic mucin while arrow heads show non secreted mucous cells. Scale bar =50 microns. Low power magnification representative images (**F**) * showing mucous plugging in the airways. Arrow heads show intracellular mucins and the arrows point to mucus plugs in the airway lumen. Scale bar= 100 microns. Graphs show box and whisker blots with min and max values on whiskers and quartile box with median value. Significant difference by unpaired T-test with * for ATP or OVA treatment difference.

**Figure 3:**
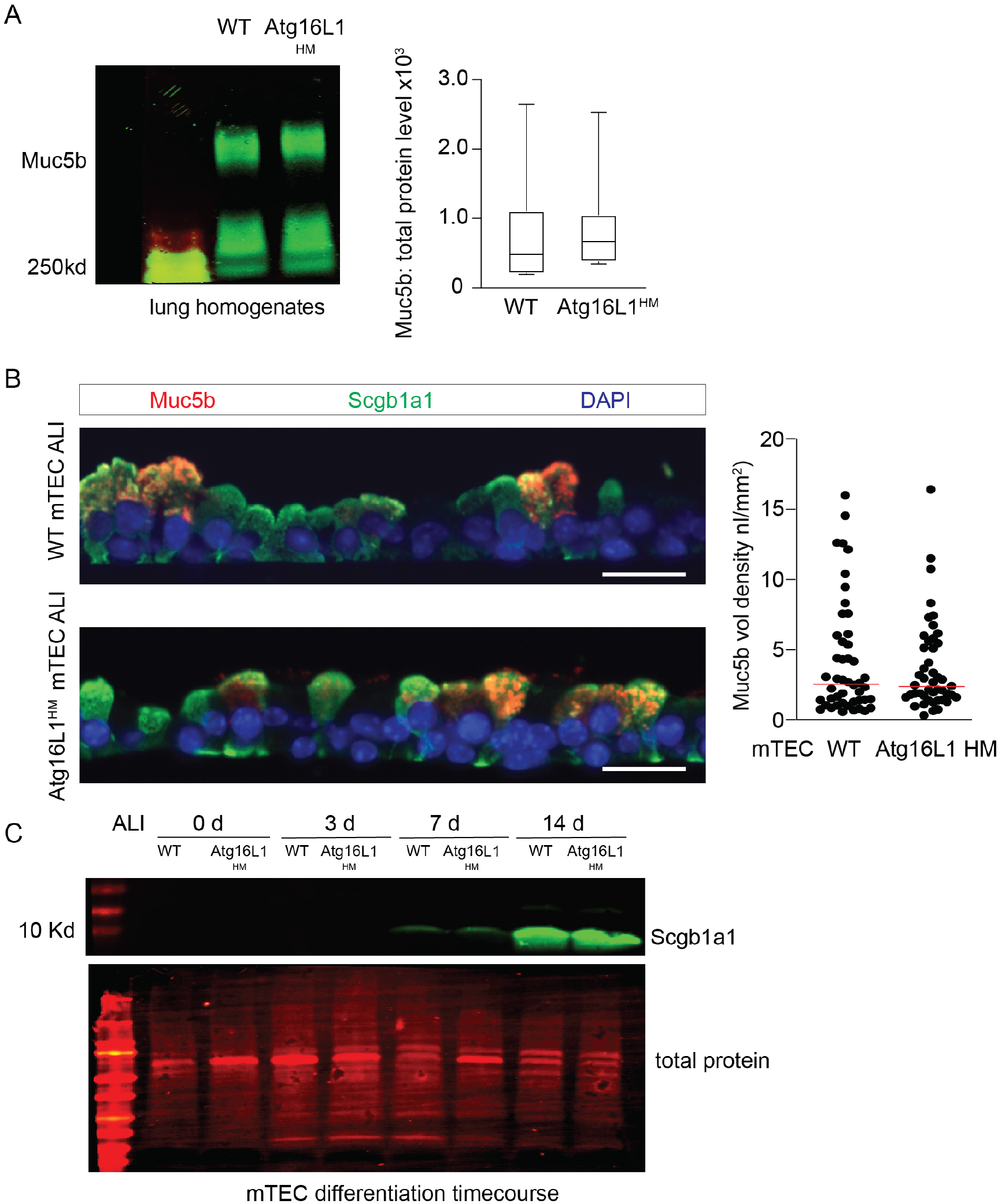
**A**) Representative immunoblots with corresponding quantification for Muc5b from lung homogenates from naïve WT and Atg16L1^HM^ mice. Values normalized to total protein levels and n=7 mice. **B**) Representative Muc5b (red) and Secretoglobin 1a1 (Scgb1a1) (green) immunostaining from WT and Atg16L1HM mouse AECs at ALI day 14 with corresponding Muc5b quantification (n=4 mTEC insert from 2 independent experiments (scale bar =25 microns). **C**) Immunoblot for Secretoglobin 1a1 (Scgb1a1) levels from WT and Atg16L1^HM^ mouse AECs at day 0, 3,7, and 14 post ALI (representative of two independent mouse AEC preps). Significant difference by unpaired T-test with * for genotype difference.

### Airway mucous cells are highly enriched for lysosomes

The autophagosome-lysosome pathway provides a means for protein degradation of proteins, aggregates(45), and organelles(5, 12, 46, 47). As lysosomes have a much longer half-life than autophagosomes(48, 49), we reasoned that if the mucin granules were being degraded in the autophagosome-lysosome, we would find abundant lysosomes in close proximity with mucin granules. Transmission electron microscopy (TEM) images from primary human airway epithelial cells (AECs) treated with IL-13 for 7d under ALI conditions revealed numerous lucent granules consistent with mucin granules(21, 50, 51) concentrated on the apical surface of mucous cells. We also observed numerous electron dense granules consistent with lysosomes(2, 52–54) that were frequently communicating directly with mucin granules (**Figure 4A**). There were also several examples of autolysosomes with remnants of the autophagosome double membrane and electron dense granules merging with mucin granules (**insets of Figure 4A**). To verify the location lysosomes in mucous cells, we next examined lysosomal membrane marker, LAMP1, immunostaining by super resolution structured illumination microscopy (SR-SIM) in human AECs treated IL-13 for 7 days or airway epithelium from mice challenged with OVA. There was significant concentration of LAMP1 positive structures closely approximating MUC5AC positive granules (**Figure 4 B and C**) in human AECs and in Muc5ac and Muc5b positive granules in OVA challenged mouse airways. We also observed a strong concentration of the lysosomal membrane marker, Lamp2A, surrounding Muc5ac granules. However, there was little staining of either Lamp1 or Lamp2A in ciliated cells (**Supplemental Figure 2 A-C)**. After a thorough review of the OVA challenged mouse airways, we observed two Lamp1 staining patterns. Larger well-formed mucous cells with abundant Muc5ac or Muc5b granules often had little Lamp1 staining while mucous cells with fewer mucin positive granules had more abundant Lamp1labeled lysosomes that were adjacent to mucin granules (**Figure 4 C-F**). This indicates that Lamp1-labeled lysosomes are not essential components of the protein exocytic machinery for secretion. Instead the variation in lysosomal staining suggests that there is a cycling population of mucous cells with distinct stages and lysosomes were abundant only in later stages. Secondly, we found frequent co-localization of Lamp1 labeled lysosomes and isolated, wayward Muc5b granules that were in a basilar pattern in the supra-nuclear cytoplasm (**Figure 4E)**. The latter Lamp1 staining pattern may be related to quality control of granules that fail to transport to the apical surface along the cytoskeletal microtubular highway. As Muc5ac is highly induced by inflammation, and then returns to baseline levels during resolution in animal models and exacerbations of human airway disease, we focused our attention on the role autophagy regulating resolution of mucous cell metaplasia by degrading Muc5ac.

**Figure 4:**
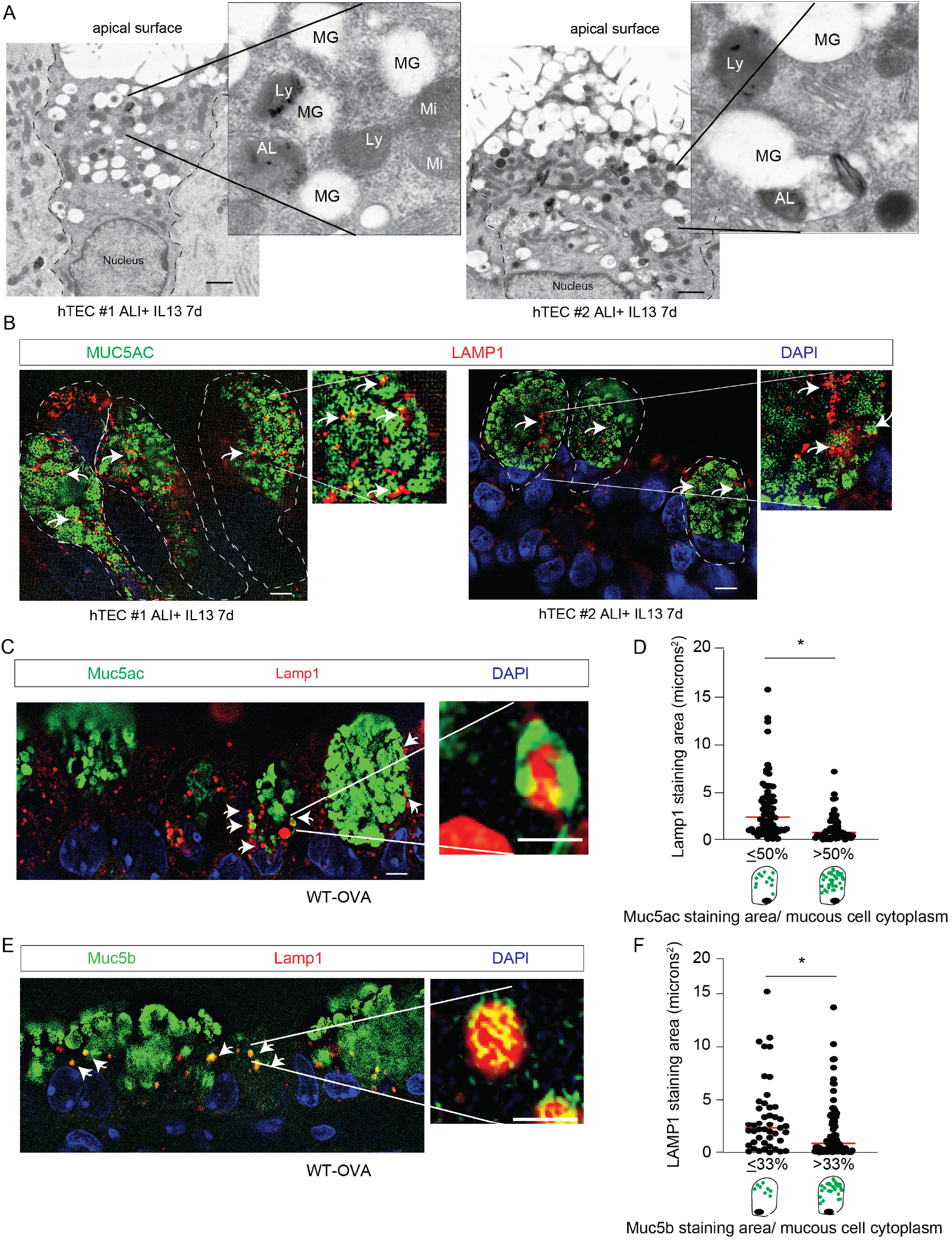
Airway mucous cells are enriched with lysosome structures and LAMP proteins. **A**) Representative from transmission electron microscopy (TEM) images from human AECs treated with 7d IL-13 (10ng/mL) under ALI conditions. Each image is from a different mucous cell from 2 unique AEC donors. Scale bar =1 micron. Ly: lysosome, MG: Mucin granule, AL: autolysosome, Mi: mitochondria. Arrows point to autophagosome membranes in the evolving autolysosome. **B**) Representative LAMP1 (red) and MUC5AC (green) immunostaining from two donors AEC donors ± IL-13 7d. Representative image of OVA challenged wildtype mouse airways with Muc5ac staining by lectin UEA1 **C**) or Muc5b **E**) (green) and Lamp1 (red). Scale bar = 5 microns. DAPI for nuclear staining. Image insets with scale bar of 1 micron. **D,F**) Quantification of Lamp1 staining area between mucous cells with ≤or > 50% Muc5ac staining area in the cytoplasm or <33 or ≥33% Muc5b staining area in the cytoplasm. N=130 mucous cells from 3 different mice. Significant difference by unpaired T-test with * for mucous cell difference.

### Persistent retention of airway Muc5ac and Muc5b in autophagy deficient mice during resolution of mucous cell metaplasia

Little is known about factors that regulate resolution of mucous cell metaplasia. It is possible that all the mucin granules are released into the airway. Alternatively, some of the granules may be degraded. Given the role of autophagy in recycling of cytoplasmic proteins, we next examined whether autophagy deficient mice had defects in resolution of mucous cell metaplasia related to reduced cytoplasmic mucin breakdown. Mice given OVA sensitization and challenge had substantial increase in Muc5ac production by qPCR that decreased by day 10 and returned to baseline by day 17 (**Figure 5A**). We previously reported that levels of autophagosome marker, LC3B II, remain elevated in AECs even after withdrawal of IL-13 or IL-4 from culture media suggesting persistent autophagy activity during resolution of mucous cell metaplasia(8). We therefore examined isolated lysosomes derived from mouse lung homogenates following OVA stimulation. These lysosomal isolates were highly enriched for LC3 II compared to the cytoplasm and levels peaked at days 0 to 3 post OVA before falling. (**Figure 5 B)**. Interestingly, while autophagosome cargo protein, SQSTM1, was much lower in the lysosome than cytoplasm, we found higher SQSTM1 levels at day 0 post OVA that then declined at day 3 to 10 post OVA. This suggests that autophagosome cargo was not being effectively degraded at peak airway inflammation but later during resolution. Next, we measured lysosome activity of cathepsins B and L which are cysteine proteases representative of lysosome enzyme activity. Cathepsin B and L proteolytic activity increased with OVA challenge but was highest days post OVA (day 3), and then declined by day 10. This indicates that lung lysosome proteolytic activity peaks during the early resolution phase. Based on our findings from naïve Atg16L1^HM^ mouse lung homogenates (**Supplemental Figure 1A-C**), we hypothesized that isolated lysosomes from autophagy deficient mice would have higher LC3 II and SQSTM1 levels. Indeed, we found Atg16L1^HM^ mice had higher levels of LC3B II and SQSTM1 levels at all time points. However, the greatest difference was day 0 (**Figure 5 D-F**). Our findings indicate that autophagy flux or proteolytic activity, is only fully consummated in the days after peak airway inflammation during resolution and autophagy deficient mice have further reduced autophagy flux.

**Figure 5:**
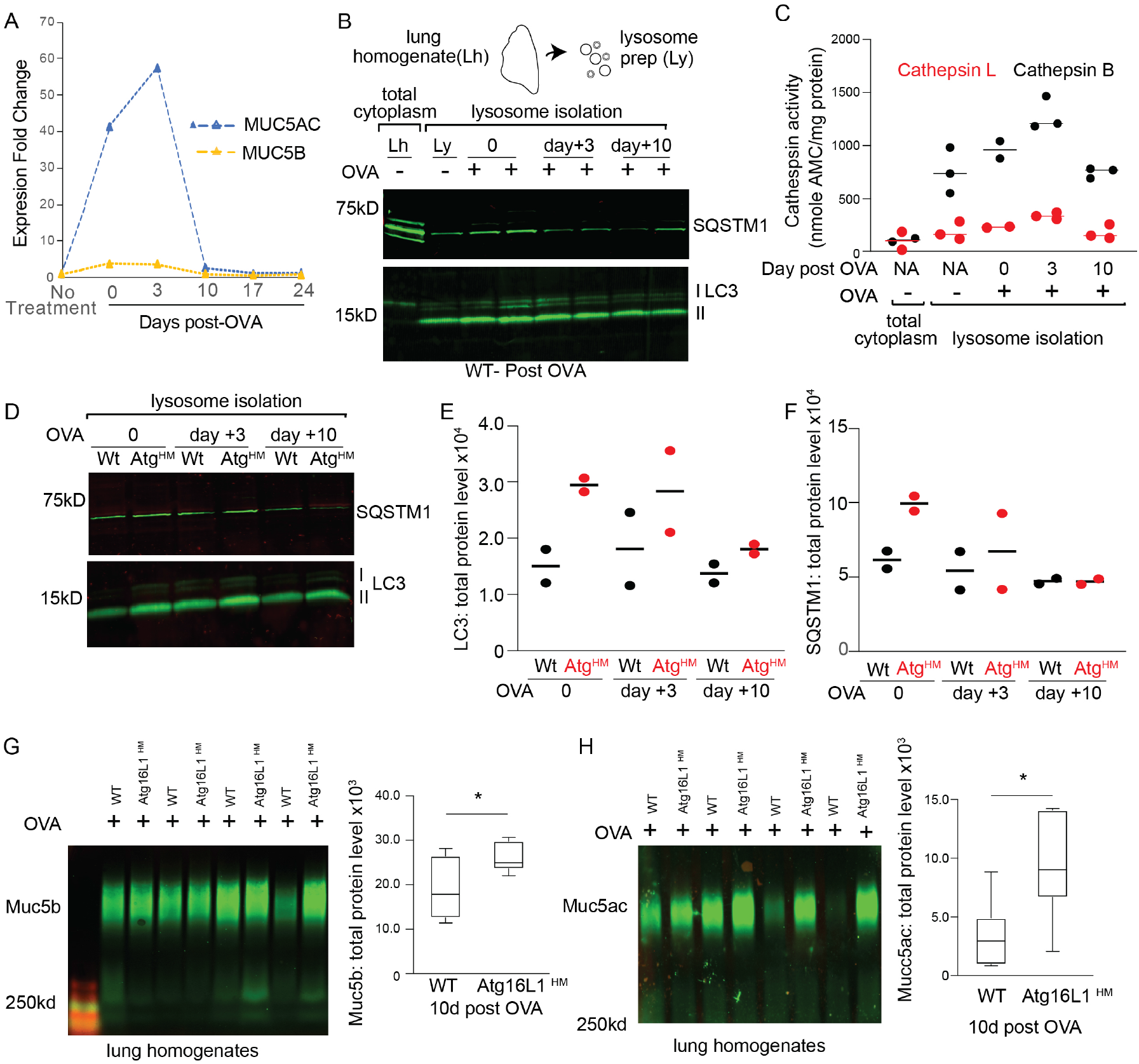
Atg16L1 deficient mice have slower resolution of cytoplasmic Muc5ac following Type 2 airway inflammation. **A**) Muc5ac and Muc5b expression were measured by qPCR in WT mouse lung homogenates in a time course experiment at 0, 3, 10, 17, and 24 days after the last OVA treatment. Data reported as fold change vs. non-treated mice. **B**) Representative LC3 immunoblot from whole lung homogenates (**Lh**) or after lysosome isolation (**Ly**) was performed from WT mouse lung homogenates ±OVA sensitization and challenge and then day 3 and 10 post OVA. (n=2 per group) **C**) Cathespin B and L proteolytic enzyme activity was measured from either lung homogenates (**Lh**) and from isolated lysosome fractions (**Ly**) from mouse lung homogenates ±OVA. Enzyme activity normalized by total protein (n=3 per group). **D**) Representative LC3 and SQSTM1 immunoblots from WT and Atg16L1HM mouse lung lysosomes at day 0,3, and 10 post OVA inflammation. Quantification of LC3 (**E**) and SQSTM1 (**F**) at day 10 post OVA from WT and Atg16L1^HM^ lysosomal isolates. (n=2 per group**). G,H**) Representative images of immunoblots of Muc5b and Muc5ac in WT and Atg16L1^HM^ mouse airways at 10 days last OVA challenge with corresponding quantification and band density values normalized to total protein levels. (n=6 mice per group). Graphs show box and whisker blots with min and max values on whiskers and quartile box with median value. Significant difference by unpaired T-test with * for genotype difference.

We then focused our attention on the impact of autophagy-mediated degradation on the remaining cytoplasmic mucins during the resolution phase at day 10 after the last OVA challenge. We found a significantly greater amount of Muc5ac and Muc5b in the airways of Atg16L1^HM^ mice (**Figure 5G-H**). A similar finding was observed in mice 10 days after 3 doses of intra-nasal IL-33 (**Supplemental Figure 3**). The return to mucin homeostasis was slowed in autophagy deficient mice with persistent elevated mucin levels. These data suggest that blocking autophagy during the resolution phase of type 2 cytokine airway inflammations models of airway mucous cell metaplasia was associated with persistent elevation in cytoplasmic mucin granules. We next sought to determine whether cytoplasmic mucins can be degraded by activating the autophagy pathway exogenously.

### Inhibition or activation of autophagy-mediated degradation regulates intracellular MUC5AC levels

Calu-3 cells, which polarize and produce MUC5AC in granules(24, 55, 56) were transfected with ATG16L1 or LC3B siRNA (**Figure 6 A-C**). We found increased intracellular MUC5AC granules by immunostaining in the ATG16L1 or LC3B deficient Calu-3 cells (**Figure 6D)**. This indicates that MUC5AC is a target of the autophagosome-lysosome protein degradation system and agrees with our earlier work that ATG5 deficient IL-13-challenged human AECs had increased cytoplasmic MUC5AC(57). We next asked if activation of autophagy by mTOR inhibition influenced intracellular MUC5AC levels. Torin1 specifically inhibits mTORC1 and TORC2 kinase activity in mammalian cells and leads to activation of autophagy(58). We have previously shown in primary airway epithelial cells that torin1 increases autophagy-mediated degradation with increased LC3 II and decreased LC3 I. Levels of LC3II increase further when autophagosome-lysosome fusion is blocked with chloroquine(8). In a similar fashion, Calu-3 cells under ALI conditions treated with torin1 for 18h had increased LC3 II and reduced LC3 I protein levels and decreased phosphorylated mTOR (**Supplemental Figure 4 A-C**). Torin1 did not lead to changes in MUC5AC gene expression, rather Torin1 reduced cytoplasmic MUC5AC protein levels by mucin immunoblotting and immunostaining. (**Figure 7 A,B**). The effect of Torin1-mediated reduction in cytoplasmic MUC5AC granules was mitigated when LC3B was depleted by siRNA (**Figure 7C**). Finally, we found examined the effect of autophagy activation by Torin1 in human AEC differentiated under ALI conditions. Again, there was decrease in MUC5AC and to a lesser extent MUC5B protein levels (**Figure 7 D,E**). These findings indicate the cytoplasmic MUC5AC can be degraded by the autophagosome lysosome pathway.

**Figure 6:**
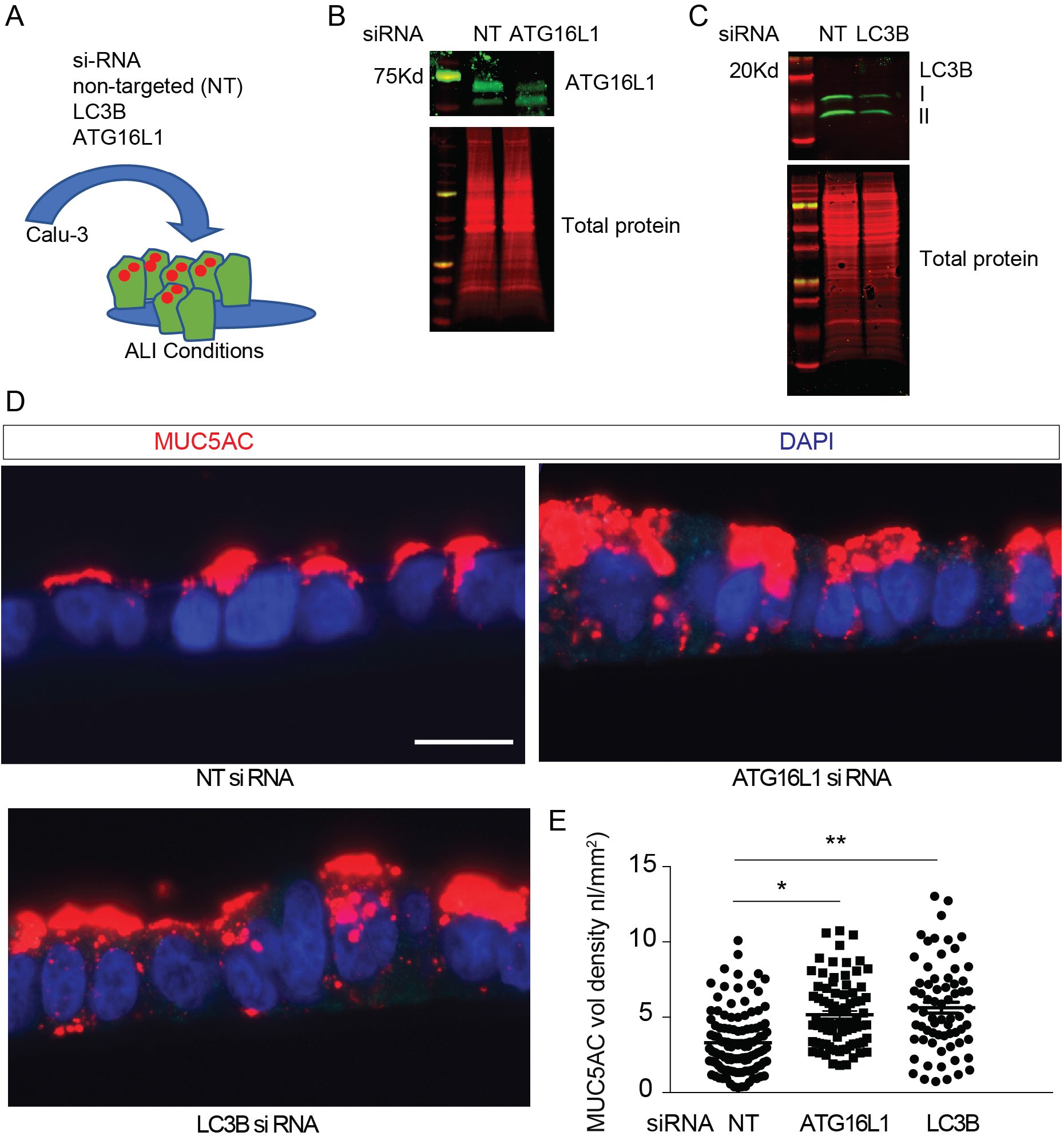
A) Autophagy deficient airway epithelial cells have increased cytoplasmic MUC5AC. Schematic for targeted deletion ATG16L1 and LC3B levels in Calu-3 cells under ALI culture conditions. Representative immunoblots for Calu-3 cells transfected with non-targeted (NT), ATG16L1siRNA (**B**) or LC3B siRNA (**C**). Band density values normalized to total protein levels. (**D**) Representative immunostaining using mouse anti-MUC5AC in Calu-3 cells transfected with non-targeted control (NT), LC3B, or ATG16L1 siRNA and corresponding quantification (**E**). Scale bar =20 microns. (n=3 experiments with 10-20 microscopic images per sample). Significant difference by ANOVA with Tukey's post-hoc comparisons are noted by * NT vs Atg16L1 siRNA treatment difference or ** for NT vs. LC3BsiRNA differences.

**Figure 7:**
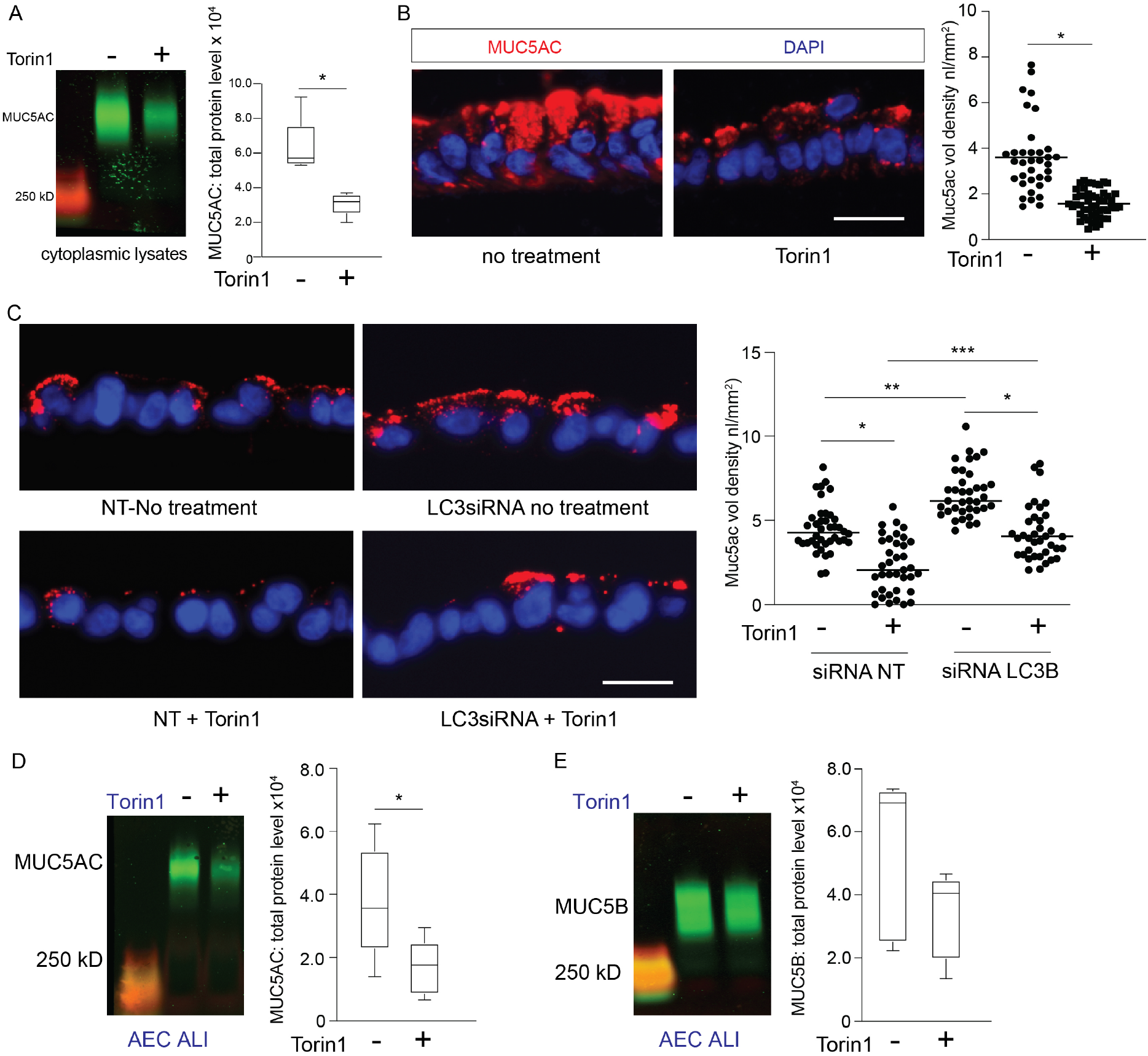
Autophagy activation leads to reduced cytoplasmic MUC5AC. **A**) Representative immunoblot of MUC5AC from cell homogenates from Calu-3 treated with torin1 (10μM) for 18hr with corresponding quantification by total protein levels (N=5). **B**) Representative MUC5AC immunostaining in Calu-3 cells under ALI conditions with torin1 treatment as described in Supplemental Figure 4 **A,B**. Scale bar =25microns. Corresponding quantification of MUC5AC staining volume density. N=3 experiments with 10-20 microscope images taken per sample. **C**) Representative MUC5AC immunostaining of Calu-3 transfected with NT or LC3B siRNA and treated ± Torin1 (10μM) for 18hr with corresponding quantification of MUC5AC volume density. N=2 experiments with 20 microscope images taken per sample. Representative immunoblot of MUC5AC **D**) and MUC5B (**E**) from cell homogenates from human non-diseased AECs under ALI conditions treated with torin1 (10μM) for 18hr with corresponding quantification by total protein levels (N=5). Significant difference by unpaired T-test with * for torin1 difference. For quantification of **E**) significant difference by ANOVA with Tukey's post-hoc comparisons are noted by * for torin1 treatment difference or ** and *** for siRNA differences.

## Discussion

Here, we show for the first time, that autophagy-mediated degradation is essential for removing excess mucin granules during mucous metaplasia and particularly during resolution. Atg16L1^HM^ deficient mice have reduced autophagy flux and accumulate increase intracellular Muc5ac granules at peak and particularly at resolution phases of mucous cell metaplasia. This difference was not due to defects in baseline or ATP-stimulated secretion. Instead, we proposed that autophagy deficient mice had reduced ability to degrade cytoplasmic mucin granules. In support of this new paradigm, we found that airway mucous cells are enriched in Lamp1-labeled lysosomes that closely approximate or engulf to mucin granules. Autophagosome marker LC3B II and degradation marker SQSTM1 isolated form lung lysosomes are highest during early the early resolution period and then decrease. This indicates that while airway inflammation increases the number of autophagosomes, autophagy flux (protein degradation) occurs most efficiently during resolution. Correspondingly, Atg16L1^HM^ deficient mice showed a delay in resolution of mucous cell metaplasia.

Inflammatory insults, such as OVA or IL-33, lead to massive increases in Muc5ac protein synthesis, ER stress(31, 59), and oxidative stress(8, 60). Our findings suggest that airway mucous cells utilize the autophagosome-lysosome protein degradation system to break down mucin proteins to restore cellular homeostasis after the inflammatory insult is over. It may seem counter intuitive for mucous cells to provide an alternative fate for secretory proteins. However, this degradation function is likely a situational phenomenon reflecting the drive to restore cellular homeostasis. As inflammation reduces, the drive to produce and secrete mucin granules decrease. To our knowledge, this is the first observation that airway secretory cells degrade cytoplasmic mucin granules by autophagy.

We, and others, have reported that autophagy is activated in the airway epithelium of T2 airway inflammation disease models(7–9). These observations are largely based on the finding of increased LC3B II in the presence of autophagosome-lysosome inhibitors by immunoblot or immunostaining from mouse lung homogenates or isolated airway epithelium. However, careful interpretation of that earlier data suggests that while the number of autophagosome structures is increased during airway inflammation, autophagosome flux or the proteolysis of the contents of the autophagosome is also, at least, partially inhibited. Indeed, our data from isolated lysosomes shows higher LC3 and SQSTM1 levels during the early resolution period that decrease over time. Our findings support a biphasic autophagy response with autophagy activation in response to external stimuli during inflammation and autophagy consummation during resolution with fusion of lysosome and proteolysis. Pathways that regulate lysosomal biogenesis, functional, and degradation (lysophagy) in the airway epithelium are not well understood. Here, we show that lysosomes are easily identified in the apical region of mucous cells and closely approximate with granules. Future studies are needed in the airway to elucidate the signals that regulate the formation, function, and degradation of lysosomes and how this relates to the resolution of mucous cell metaplasia.

Previous studies using chloroquine have postulated that autophagy inhibition may be a beneficial strategy in muco-obstructive airway disease(9). Our data is a caution that autophagy action is cell and context dependent. Strategies to inhibit autophagy action in the airway may have unintended consequences to prolong mucous cell metaplasia and prevent resolution. In fact, our data shows that activation of autophagy through mTOR inhibition lead to reduced levels of MUC5AC. Previously we have shown that AECs deficient in ATG5 and ATG14L1 had reduced IL-13-mediated apical secretion.(7) This finding agreed with similar data by collaborators in the gut epithelium, showing autophagy proteins are required for Muc2 secretion in the colon based on a ROS-mediated mechanism.(61, 62) Airway mucin secretion relies on a baseline and stimulated secretion with distinct exocytic machinery. Our current findings examine both baseline and ATP-stimulated secretion and find no relationship with Atg16L1 deficiency. In the present study, we did not investigate ROS-mediated mucin secretion which is less well understood. It is possible that autophagy regulatory proteins in a non-canonical fashion (not related to autophagy-lysosomal degradation) regulate mucin secretion when inflammation (ROS) is highest while canonical autophagy regulates mucin degradation as a means to restore cellular homeostasis during resolution.

This study provides evidence to alter the existing paradigm that only production and secretion regulate the level of cytoplasmic mucin content. We propose that there is a third factor, degradation, that plays a role in resolving mucous cell metaplasia by degrading mucin granules by autophagosome-lysosome action. Our findings offer a new potential therapeutic strategy to speed the resolution of mucous cell metaplasia in those with exacerbations of muco-obstructive lung diseases.

## Materials and Methods

### Mice

C57BL/6J wild-type (WT) were purchased from The Jackson Laboratory (Bar Harbor, ME). Atg16 hypomorph (HM) mice with global deficiency of Atg16L1 gene on the same C57BL/6J background(43) were obtained courtesy of Thaddeus Stappenbeck (Washington University, St Louis, MO). Male and female mice were utilized in equal numbers between 12-16 weeks of age for all experimental studies. Food and water were provided *ad libitum*.

WT or Atg16L1^HM^ mice were lightly anesthetized by isoflurane inhalation before intranasal inhalation of 50 μl of sterile saline or 1 μg of IL-33 daily for 5 days. Alternatively, mice were OVA-albumin sensitized by intraperitoneal injection x3 over 7 days. After 7-day incubation period, mice were subsequently challenged by nebulization daily over 5 days as previously described(8). To assess the role of Atg16L1 in the secretory response of intracellular mucins to ATP, a subset of WT and Atg16L1^HM^ mice were placed in a plexiglass chamber and exposed to nebulized ATP (100 mM diluted in sterile water) 30-60 minutes prior to sacrificing the animal as previously described(25, 63).

### Cell culture techniques

Mouse tracheal epithelial cells (mouse AECs) were isolated after pronase (Sigma-Aldrich, 9036-06-0) digestion of WT and Atg16L1^HM^ mouse tracheas and grown on supported membrane inserts (Corning #3460 and #3470) as previously described(64–66). Mouse AECs were expanded (5-8 days) using DCI media containing: Isobutyl methylxanthine (IBMX), final concentration 0.1 mM (Sigma, I5879), 8-Bromoadenosine 3',5'-cyclic monophosphate sodium (8-BAMS), final concentration 0.1mM (Sigma, B7880), Calcium chloride final concentration 1mM (Sigma), L-glutamine final concentration in Advanced DMEM/F12 500 ml (Gibco#12634-010) and then fed with PneumaCult ALI media (Stemcell Technologies # Catalog #05001) under air liquid interface (ALI) conditions. Non-diseased human airway epithelial cells (AECs) were derived for culture from excess airway tissue donated for lung transplantation as previously described(7). COPD cells were derived from the large airways after lung transplantation. After 5-7 days of expansion with BEGM media (Lonza #CC-3170), air-liquid interface (ALI) conditions were established and cells were fed with PneumaCult ALI media. Calu-3 epithelial cells were chosen for their ease in transfection and ability to polarize under ALI conditions and secrete mucin, MUC5AC apically(55, 56). Calu-3 cells obtained from ATCC (#HTB-55) between passage 25-35, were cultured in Eagles Minimal Essential Media (ATCC #30-2003) media plus 10% FBS on supported membranes as above for AECs. Cells were utilized 7-14 days after the establishment of ALI conditions with cell confluency. ATCC authenticated the identification of Calu-3.

### Immunoblotting

Cell lysates cultured from Calu-3 cells or mouse AECs were isolated in RIPA buffer with protease inhibitor on ice for 20 minutes with brief water bath sonication. Cell lysates for mTOR and P-mTOR detection were lysed in a NP-40 buffer: 50mM HEPES, 0.5% NP40, 2.5mM EDTA, 50mM NACL, 10mM sodium pyrophosphate, 50mM sodium fluoride, and protease inhibitor cocktail. Samples underwent electrophoresis on a gradient 5-14% gradient acrylamide gel and then were transferred on PDVF membranes over 2 hours. The blot was rinsed with TBS with 0.1% tween and stained with REVERT Total Protein Stain with conjugated 700nm fluorochrome (LI-COR, Lincoln, NE) for 5 minutes. Blots were imaged using an Odyssey CLx Imager (LI-COR) for total protein detection. Membranes were blocked with either 5% BSA in TBS with 0.1% tween for or 5% non-fat milk for one hour at room temperature and then incubated with primary antibody overnight at 4°C in the diluted appropriate blocking buffer: Scgb1a1 1:500 vol;vol, (Seven Hills Bioreagents #WRAB-3950), Atg16L1 1:1000 vol;vol (Cell Signaling #D6D5), total mTOR and phosphorylated (Serine 2448) mTOR (1:1000 vol;vol (Cell Signaling #2972S and #2971S). After washing with TBS and 0.1% tween, the IRDye 800nm CW goat anti-rabbit (Li-Cor 926-32211) at a 1:10,000 dilution in blocking buffer was added to the blot and incubated at room temperature for 1 hour, and levels were normalized to total protein levels. Calu-3 total cell lysates were solubilized for LC3 immunoblot and run on a 15% polyacrylamide gel for transfer onto PDFM membranes as previously described(7, 8). LC3 was detected by rabbit polyclonal LC3B II 1:500 vol;vol (Sigma-Aldrich L7543). Levels were normalized either to ACTIN or total protein levels.

Mouse lungs were prepared for immunoblotting as previously described(8). Briefly following euthanasia, the right heart was cannulated, and sterile PBS with heparin sulfate (0.67Units/mL) was infused to remove blood from the pulmonary vasculature. Lungs were homogenized RIPA buffer with protease inhibitor using a tissue dissociator (GentleMacs Dissociator Miltenyi Biotec). Homogenates were then centrifuged at 10,000 rpm and 4°C and the supernatants were collected and prepared for either Lc3 or Atg16L1 detection as described above.

### Mucin immunoblots

After euthanasia, excised mouse lungs were snap frozen in liquid nitrogen. Lungs were then later homogenized in PBS with protease inhibitors (Roche, Mannheim, Germany) using a gentleMACS dissociator (Miltenyi Biotec GmbH, Bergisch Gladbach, Germany). Subsequently the lysates were centrifuged for 30 min at 16,000 rpm at 4°C. Lung homogenate supernatants were then collected and quantified. Loading buffer (containing 5 mM dithiothreitol (DTT) for Muc5b detection or no DTT for Muc5ac by UEA-1 detection was used. Electrophoresis was performed on a 0.8% agarose gel. The gel was incubated in 10mM DTT saline sodium citrate (SCC) buffer (3.0 M sodium chloride, 0.3 M sodium citrate, pH 7.0) at room temperature for 20 minutes then rinsed in water for Muc5b detection. Gels with Muc5ac by UEA-1 detection were rinsed with SCC buffer without DTT and proceeded directly to transfer. The protein was then transferred onto nitrocellulose membrane (BioRad, Germany) using a vacuum Blotter (Model 785, BioRad). Blots were rinsed with TBS and blocked with TBS with 5% non-fat milk for 1 hour at room temperature and then incubated with mouse monoclonal antibody for Muc5b (1:500 vol:vol) or lectin UEA-1 L8146 1mg/mL (1:1000 vol:vol) (Sigma-Aldrich) overnight at 4°C. IRDye 800nm CW Goat anti-Mouse (Li-Cor) at a 1:10,000 dilution for Muc5b or rabbit anti-lectin UEA-1 (Sigma, U4754) at a 1:1,000 dilution in blocking buffer for Muc5ac was added to the mucin blot and incubated at room temperature for 1 hour. IRDye 800nm CW goat anti-rabbit (Li-Cor) at a 1:10,000 dilution in blocking buffer was added to the Muc5ac blot and incubated at room temperature for 1 hour. Mucin blots were normalized for quantification using total protein levels as previously described(63).

### Assessment airway epithelial immunostaining

Following euthanasia and lung perfusion, the whole lungs were then excised and slowly inflated with 10% formalin and hung under 20cm of H20 pressure for 24 hours. Fixed lung tissue was embedded in paraffin and 5-micron lung sections were cut. Antibody and lectin-specific staining and quantification was performed on embedded lung tissue slides as previously described(63). Staining for lysosomal membrane markers, LAMP1 and LAMP2A was done with a rabbit polyclonal antibodies 1:500 vol;vol (Cell Signaling; #D2D11 and ThermoFischer #51-2200 respectively). Staining for cilia was performed using mouse monoclonal antibody to acetylated alpha tubulin 1:1000 vol;vol (Cell Signaling #11H10). Secondary fluorophore-labeled donkey secondary, species-specific antibodies were Alexa Fluor 488 or 555 (Life Technologies; #A-31570, #A21202, #A21206, #A31572). Nuclear counterstaining of DNA was obtained with 4′,6 diamidino-2-phenylindole (DAPI). Mouse airway or cultured AECs mucin volume density was quantified as described previously(8, 25). Images were obtained using a Zeiss Axio Observer Z.1 microscope and analyzed by Zen software (Zeiss). Super resolution images were obtained using a Zeiss Axio Observer Z.1 microscope and analyzed by Zen software (Zeiss). Super resolution Structured Illumination Microscopy for Optical Section (SR-SIM) obtained images with 0.110-micron Z-stack slices by Zeiss PS.1 confocal microscope. For quantification of SR-SIM images of mucous cells and lysosomal markers, a single Z-stack was chosen that contained multiple mucous cells. A common image threshold was utilized for each fluorescent channel by imageJ software. The amount of Lamp1 staining was measured for each mucous cell and dichotomized for mucous cells with ≤ or >50% Muc5ac staining area of the total mucous cell cytoplasm or ≤ or >33% of Muc5b staining area of the total mucous cell cytoplasm.

Mouse/human airway epithelial cells (AECs) and Calu-3 staining was performed in cross sections of paraffin embedded formalin inserts. First, AECs were embedded in warmed 37°C 1% agarose. After cooling, the inserts were fixed with 10% formalin and embedding in paraffin. Cross sections from the inserts were then cut and fixed on slides for immunostaining as previously described(63). Using the same antigen retrieval and blocking method described above for mouse lung sections, AECs were stained with (1:500 vol:vol dilution) mouse monoclonal antibody to MUC5AC (Thermo Fischer Scientific #MA5-12178) for human AECs and Calu3 cells. Mouse AECs were stained for Muc5b by mouse monoclonal antibody 3AE (27) and rabbit antibody Scgb1a1 (Seven Hills BioReagents WRAB-3950).

### Electron microscopy

Cultured human AECs under ALI conditions treated with 7d IL-13 (10ng/mL) were fixed with a solution containing 4% paraformaldehyde and 0.5% glutaraldehyde and processed for transmission electron microscopy as previously described(7).

### Lysosomal isolation and enzymes activity

Mouse lungs were perfused with PBS and heparin as described above. Then one lung was collected for lysosome isolation (Minute Lysosome Isolation Kit; Invent Technologies; LY034) per manufactures protocol. Briefly, lungs were dissociated by forcing the tissues again the collection column for one minute and then incubated on ice for 5 minutes. A series of column centrifugations was used to enrich lysosomes from the crude cytoplasm. The final pellet lysed in 40μl of denaturing protein SDS solubilization reagent (Invent Technologies; WA-009) and then quantified prior to immunoblot or protease activity assay. To determine lysosome protease activity from lysosomes of lungs challenged with OVA, Cathepsin activities were measured after incubating isolated lysosomes with cathepsin B and cathepsin L fluorogenic substrates as described earlier(67, 68).

### RNA interference

ATG16L1, LC3B, or control, non-targeted (NT) siRNA sequences (Dharmacon On-Target Plus; #L-021033-01,# L-012846-00-0005, #D-001810-10-20) were transfected in Calu-3 on day 0 of culture on insets. The siRNA oligonucleotides (80 nM) were suspended in Opti-MEM 1 (25%; Life Technologies, #31985-070) and Lipofectamine RNAiMAX (Invitrogen; #13778-150) as previously described(8). Depletion of ATGL16L1 and LC3B was verified on ALI 7 days by immunoblot analysis using antibodies to rabbit anti-ATG16L1 (1:1000 vol;vol ATG16L1 1:1000 vol;vol (Cell Signaling D6D5) or rabbit anti-LC3B (1:500 vol;vol Sigma-Aldrich L7543). Cells were then agarose embedded and formalin fixed for staining as described above or homogenates collected for mucin blot at ALI day 10.

### Autophagy induction and measurement of MUC5AC and MUC5B in Calu-3 and human AECs

To determine intracellular mucin levels *in vitro*, the apical surface was rinsed with 100 μl pre-warmed PBS lacking calcium and magnesium for 15 minutes at 37oc a total of three times prior to the experiment. For autophagy induction assays, cells were treated with 10μM torin1 (Sigma Aldrich #475991) and or vehicle control DMSO 1/1000 working dilution in the basolateral media and apical PBS. For mucin collection, the AECs were flash frozen in liquid N_2_ and stored at −80°C. After thawing, cells were collected in PBS with protease inhibitor, homogenized using probe sonication for 3 seconds, centrifuged for 30 minutes at 16,000 g at 4°C, and quantified. Immunoblotting was then continued as described above for mouse lung homogenates with mouse monoclonal anti-MUC5AC (Thermo Fischer Scientific #MA5-12178) and rabbit polyclonal anti-MUC5B (Santa Cruz SC-20119) (1:500 vol;vol) used respectively.

### PCR gene expression

RNA was isolated from whole mouse lung tissue using the RNeasy Spin column kit (Qiagen #75144) then reverse transcribed using the cDNA Reverse Transcription Kit (Applied Biosystems #4308228) as previously described(63). Gene expression was reported as fold change over WT-saline condition normalized to GAPDH in mouse lung tissues and calculated using the double delta CT (2ΔΔCT) method as previously described(7, 8, 63).

### Statistical Analysis

Significant difference by ANOVA with Tukey's post-hoc comparisons are noted by * for IL-33 treatment difference or ** for mouse genotype differences. Significant difference by unpaired T-test with * for treatment difference.

### Study approval

All animal procedures were approved by the Institutional Animal Care and Use Committee of the University of Nebraska Medical Center. Human AECs derived from airways of lungs not suitable for transplant or from COPD lung explants after transplant were isolated under protocols approved by the University of Nebraska Medical Center and Washington University School of Medicine.

## Author Contributions

JD Dickinson performed experiments, interpreted the data, and wrote the manuscript. K Kudrna performed experiments and helped with editing the manuscript. JM Sweeter performed experiments, assisted with data interpretation, and helped with editing the manuscript. K Hunt assisted with experiments using torin1 on Calu3 cells. P Thomes assisted with lysosomal isolation and cathepsin enzyme assays. BF Dickey assisted with experimental design and provided Muc5b monoclonal antibody. He also assisted with editing the manuscript. SL Brody assisted with experimental design, data interpretation, and manuscript editing.

## Acknowledgements

Dr. Thaddeus Stappenbeck kindly provided the Atg16L1HM mice used in this work. The UNMC Advanced Microscopy Core Facility provided assistance with image acquisition and analysis for structured illumination microscopy. Dr. Wandy Beatty from Washington University School of Medicine provided assistance with fixation and electron microscopy image acquisition of airway mucous cells. Dr. Kristina Bailey provided non-diseased and COPD derived human AECs from IRB approved tissue bank at UNMC.

Grant award acknowledgements: K08HL131992-JDD, ------BFD, -------SLB

