## Supplemental Figures for "Mucin granules are degraded in the autophagosome-lysosome pathway as a means of resolving airway mucous cell metaplasia"

**Figure Legend:**

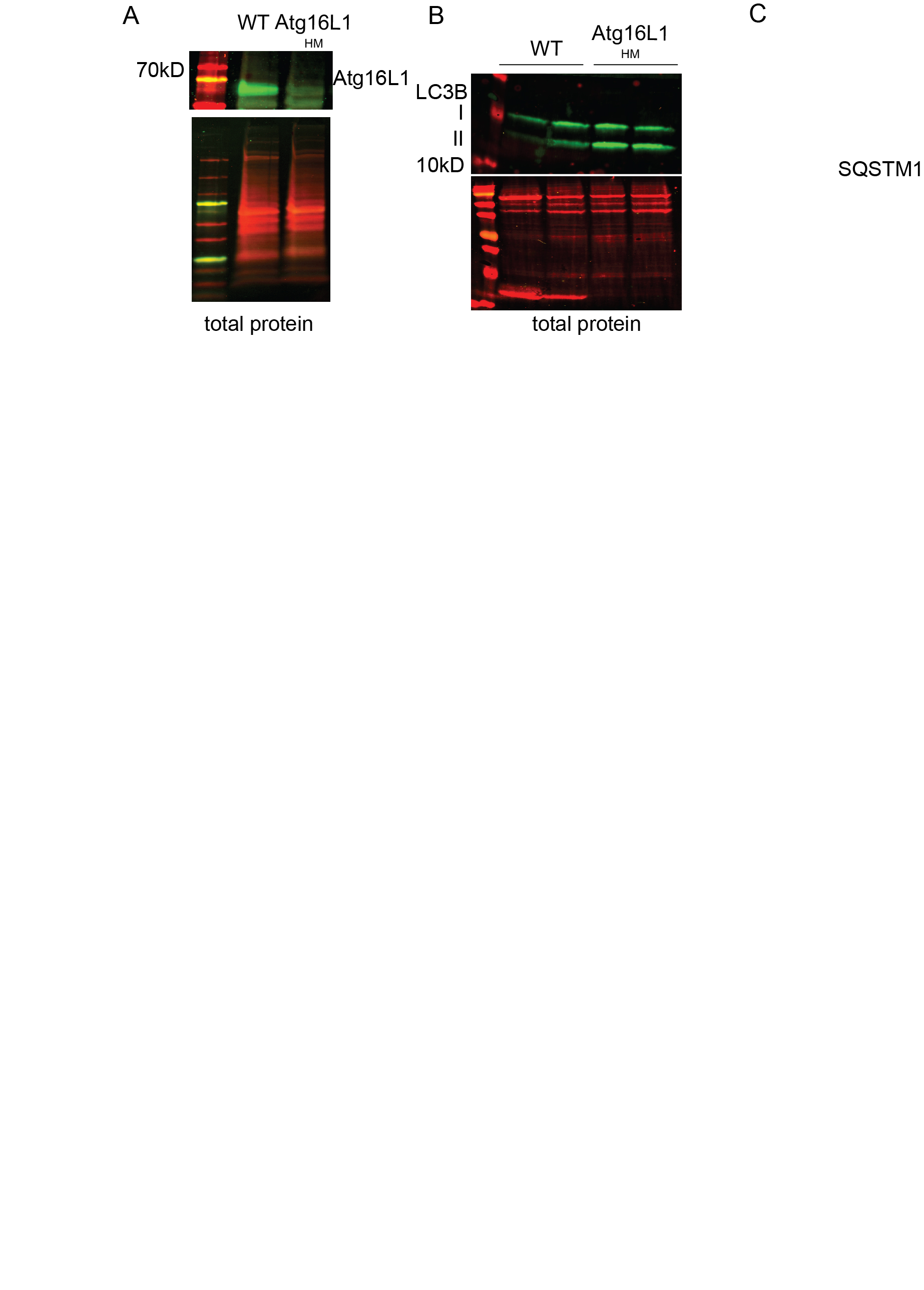

**Supplemental Figure 1**: **Atg16L1HM mice have reduced Atg16L1 and increased LC3II levels in lung homogenates.**  **A**) Representative immunoblots for Atg16L1 and LC3B **B)** levels from lung homogenates in naïve WT and Atg16L1^HN^ mice (N=4). Values normalized by total protein levels.

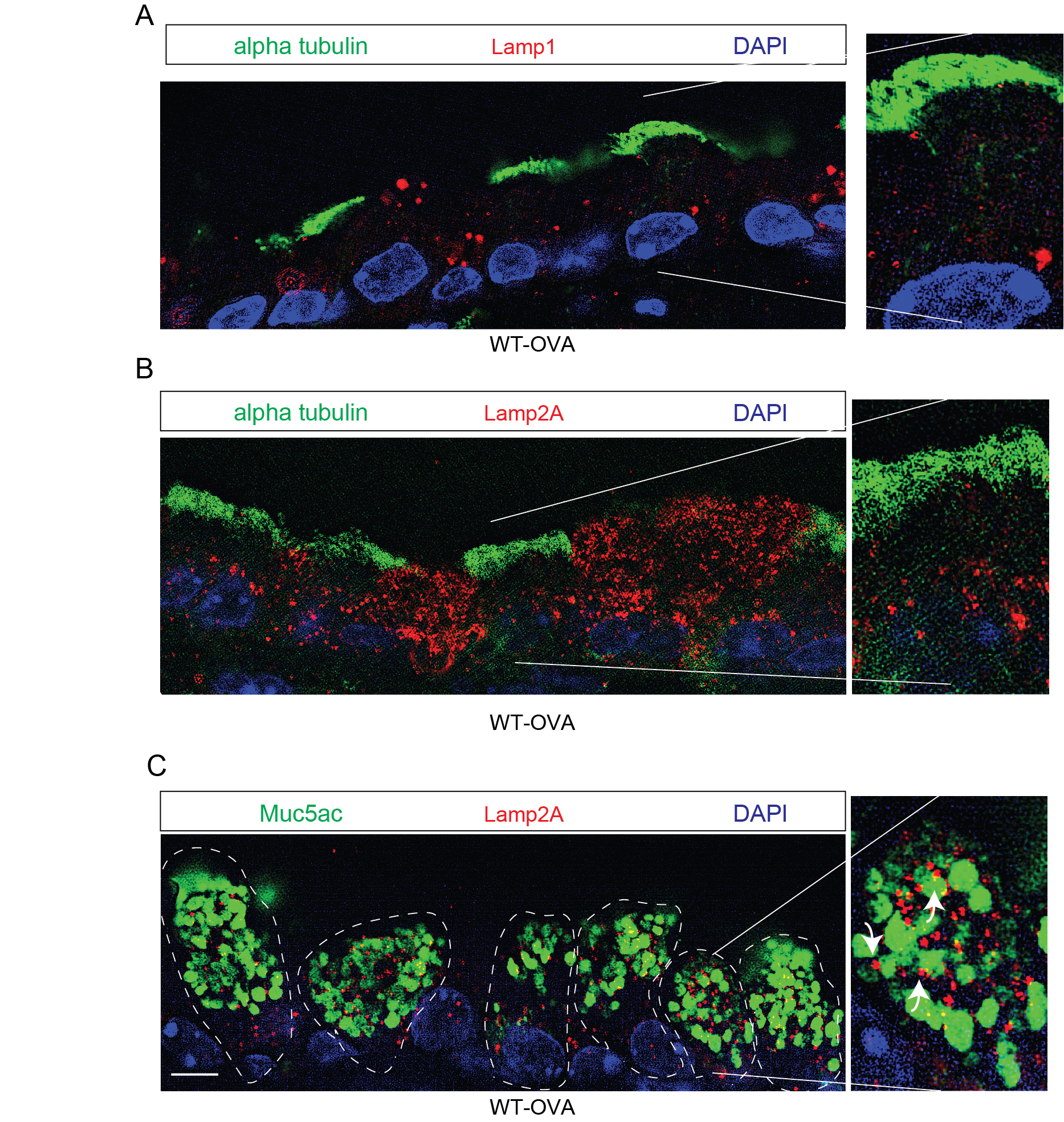
**Supplemental Figure 2**: **Lysosomal membrane LAMP proteins concentrate in the cytoplasm surrounding mucin granules of secretory cells.** **A**) Lamp1 (red), acetylated (AC) alpha tubulin (green), immunostaining in OVA challenged mice. **B**) Representative Lamp2A (red), acetylated alpha tubulin (green) or Muc5ac by lectin uea1 (green) staining in OVA challenged mouse airways. challenged mice. Scale bar =5 microns. DAPI for nuclear counter staining. Arrows in inset point to Lamp1 and Lamp2A adjacent to or overlapping with Muc5ac granules.

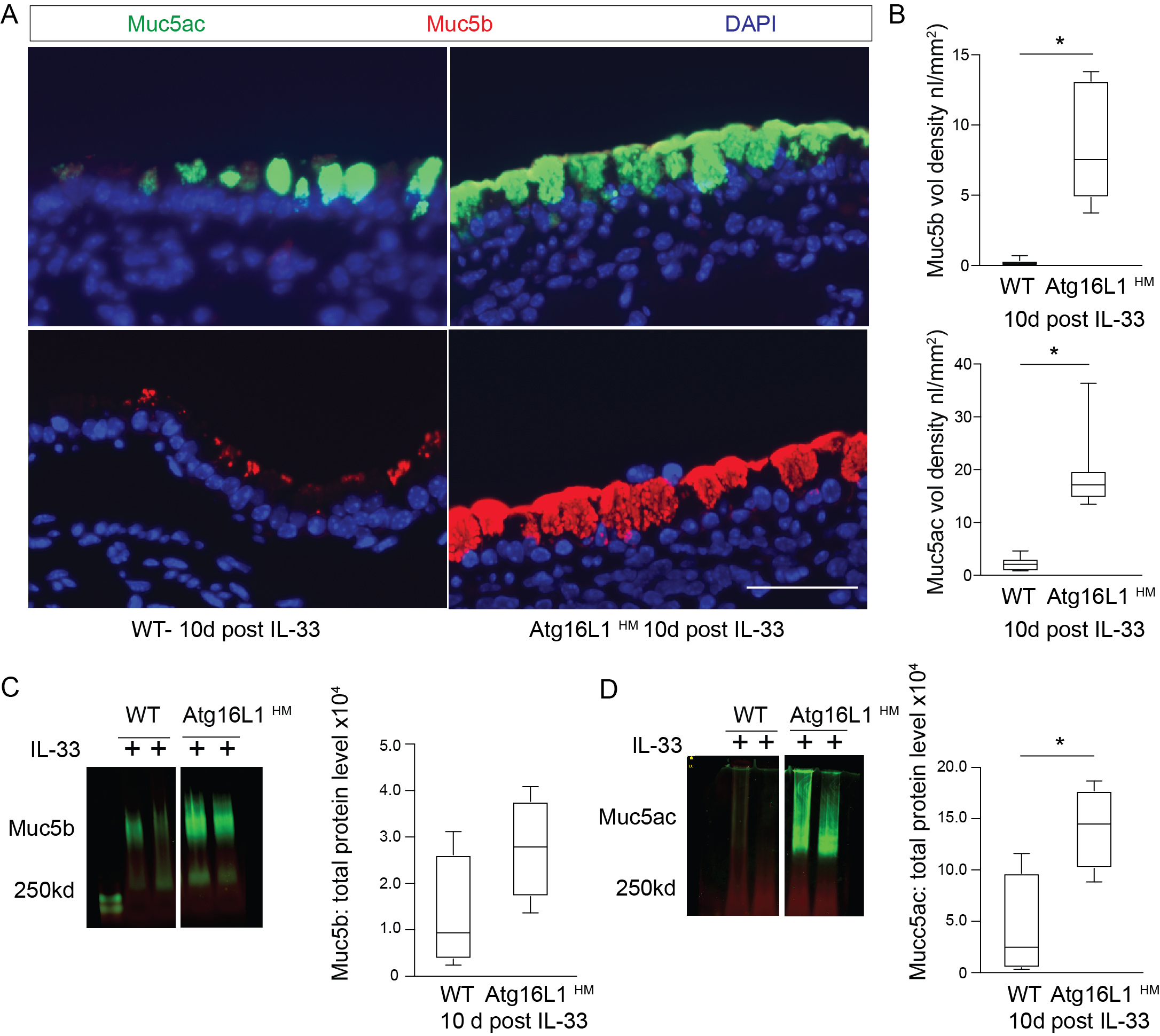

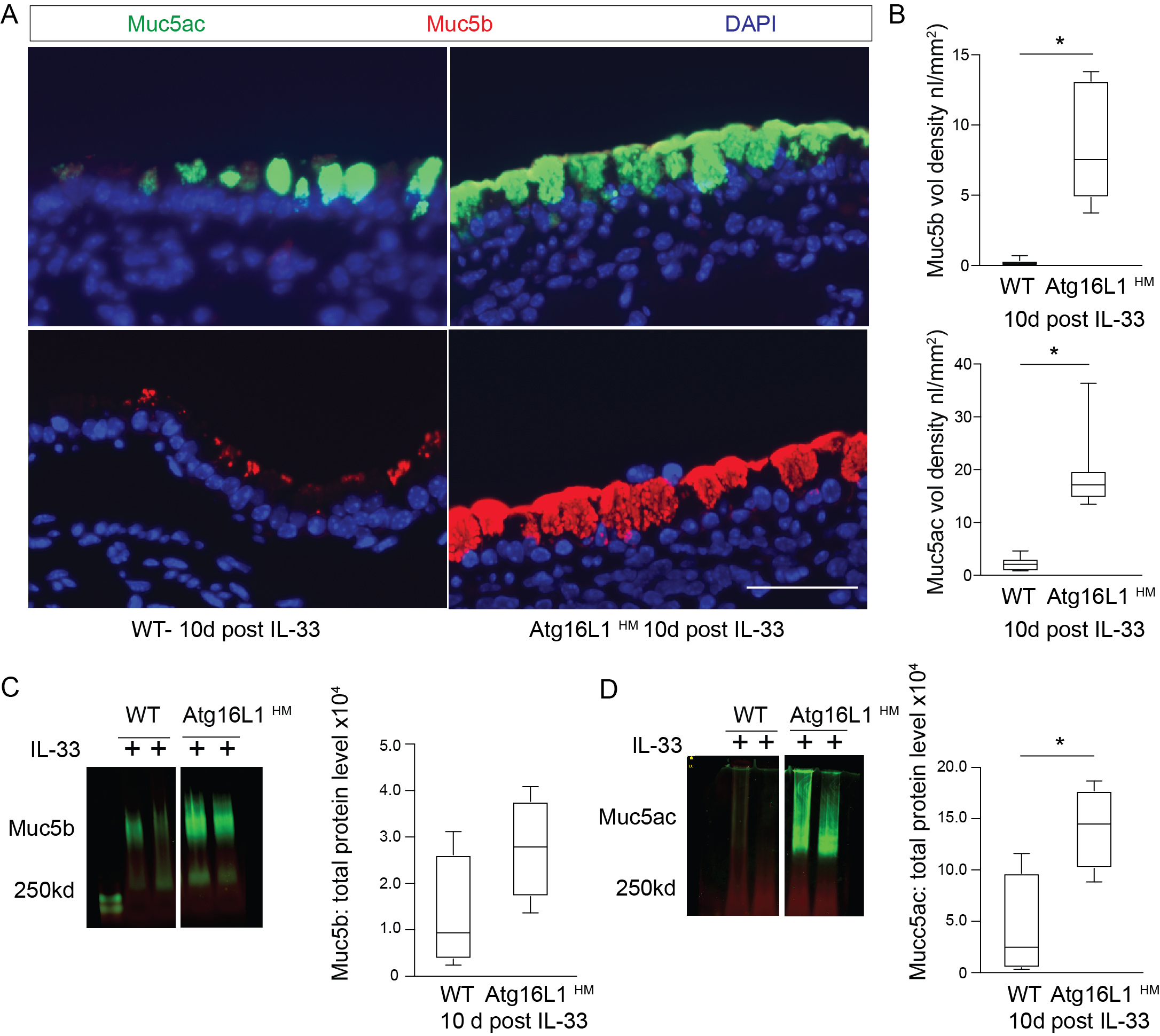

Supplemental Figure 3: **A,B**) **Atg16L1 deficient mice have slower resolution of cytoplasmic Muc5ac following IL-33 airway inflammation.** Representative immunostaining and quantification for Muc5ac and Muc5b 10 days following last of 3 challenges of intra-nasal IL-33. Scale bar equals 20 microns. **C,D**) Representative images of immunoblots of Muc5b and Muc5ac in WT and Atg16L1^HM^ mouse airways at 10 days last IL-33 challenge with corresponding quantification and band density values normalized to total protein levels (n=4 mice per group). Graphs show box and whisker blots with min and max values on whiskers and quartile box with median value. Significant difference by unpaired T-test with * for genotype difference.

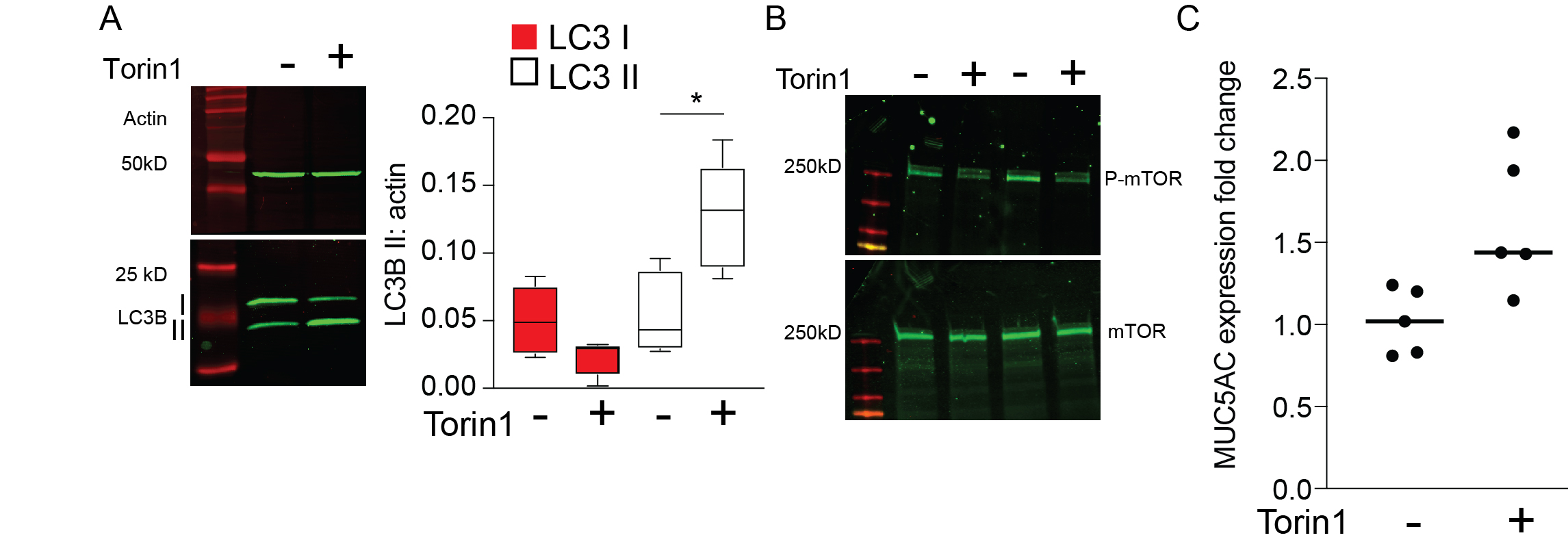
**Supplemental Figure 4**: **A**) Representative LC3B immunoblot from Calu-3 cells under ALI conditions given torin1 (10μM) for 18hr. LC3B I and II levels were each normalized to total protein. N=5 experiments. **B)** Representative immunoblot for mTOR and phosphorylated mTOR (S2448) with and without torin1 (N=5 experiments). **C**) MUC5AC expression from Calu-3 ±torin1 (10mM) 18hr. Expression normalized by OAZ1 and reported as fold change over untreated (N=5).
